# Developmental Coordination Disorder: What Can We Learn From Recombinant Inbred Mice Using Motor Learning Tasks and Quantitative Trait Locus Analysis

**DOI:** 10.1101/2022.05.18.492177

**Authors:** Kamaldeep Gill, Jeffy Rajan Soundara Rajan, Eric Chow, David G. Ashbrook, Robert W. Williams, Jill G. Zwicker, Daniel Goldowitz

**Affiliations:** Rehabilitation Sciences, University of British Columbia, Vancouver, Canada; British Columbia Children’s Hospital Research Institute, Vancouver, Canada; Department of Medical Genetics, University of British Columbia, Vancouver, Canada; Centre for Molecular Medicine and Therapeutics, University of British Columbia, Vancouver, Canada; Department of Genetics, Genomics and Informatics, University of Tennessee Health Science Center, Memphis, TN 38163, USA; Department of Occupational Science & Occupational Therapy, University of British Columbia, Vancouver, Canada; Department of Pediatrics, University of British Columbia, Vancouver, Canada; Sunny Hill Health Centre at BC Children’s Hospital, Vancouver, Canada

**Keywords:** developmental coordination disorder, motor learning, neurodevelopment, accelerated rotarod, horizontal rung, complex wheel, skilled reaching

## Abstract

Developmental Coordination Disorder (DCD) is a motor skills disorder that affects 5-6% of all school-aged children. There is an indication that DCD has an underlying genetic component due to its high heritability. Therefore, we we have explored the use of a recombinant inbred family of mice known as the BXD panel to understand the genetic basis of complex traits (i.e., motor learning) through identification of Quantitative Trait Loci (QTLs). The overall aim of this study was to utilize the QTL approach to evaluate the genome-to-phenome correlation in BXD strains of mice in order to to better understanding the human presentation of DCD. Results in this current study indicate there is a spectrum of motor learning in the pre-selected BXD strains of mice with a spectrum between high and low learning capabilities. Five lines – BXD15, BXD27, BXD28, BXD75, and BXD86 – exhibited the most DCD-like phenotype, when compared to other BXD lines of interest. The results indicate that BXD15 and BXD75 struggled primarily with gross motor skills, BXD28 primarily had difficulties with fine motor skills, and BXD27 and BXD28 lines struggled with both fine and gross motor skills. The functional roles of significant QTL genes were assessed in relation to DCD-like behavior. Only *Rab3a* (Ras-related protein Rab-3A) emerged as a best candidate gene for the horizontal ladder rung task. This gene is found to be associated with brain and skeletal muscle development. This is the first study to specifically examine the genetic linkage of DCD using BXD lines of mice.

## Introduction

Developmental Coordination Disorder (DCD) is a motor skills disorder that affects 5-6% of all school-aged children (Zwicker et al., 2012). This neurodevelopmental disorder is characterized by marked impairments in motor coordination and motor learning that significantly interferes with activities of daily activities and academic achievement (Zwicker et al., 2012). The criteria for diagnosis of DCD includes: (1) learning and execution of coordinated motor skills that are below the expected level for age and opportunity for learning; (2) motor skill difficulties significantly interfere with activities of daily living and impact academic progress, prevocational and vocational activities, leisure, and play; (3) onset is in the early developmental period; and (4) motor skill difficulties are not better explained by intellectual disability, visual impairment, and or any other neurological condition that affects movement (American Psychiatric Association, 2013). As a result of motor difficulties, children with DCD struggle with everyday tasks, such as tying shoelaces, using utensils, handwriting, physical education, and participation in leisure activities (Zwicker et al., 2012). It was once believed that children with DCD would outgrow their motor difficulties; however, recent evidence suggests that difficulties persist into adolescence and adulthood (Kirby et al., 2014).

The functional difficulties experienced by children with DCD can be attributed primarily to motor impairments in three main areas: poor postural control, poor motor coordination, and difficulty in learning motor skills (Biotteau et al., 2016; Zwicker et al., 2009). In addition to motor learning challenges, children with DCD struggle with planning motor movement, adapting to changes in external task demands, automatizing motor patterns, and generalizing motor skills in different contexts (Biotteau et al., 2016; Zwicker et al., 2009).

The cause of DCD is currently unknown; however, a large body of evidence strongly suggests that the cerebellum and/or its network of connections is a logical source of the dysfunction (Zwicker et al., 2009). The cerebellum’s key role in co-ordinated behaviours, postural control, and motor learning point to this structure underlying DCD. In support this hypothesis, Gill et al. (2020) and Zwicker et al. (2011) found structural and functional abnormalities in the cerebellum of children with DCD. Additionally, there is an indication that DCD has an underlying genetic component due to its high heritability. Lichtenstein et al. (2010) found heritability estimates of 70% for DCD using a large population-based study of 16,858 Swedish twins aged 9-12 years and validated parent telephone interviews assessing various child neuropsychiatric problems. Lichtenstein et al. (2010) also found higher concordance rates for monozygotic twins compared to dizygotic twins, indicating that there are genetic variants contributing to DCD. Despite knowing this, very little research has explored the genetic underpinning of DCD.

Although mouse models cannot replicate a human disorder, the fundamental symptoms can be approximated for the purpose of investigating the phenotypic variation and genetic causes of the motor impairments. Most animal models are developed using single or few inbred backgrounds, with little or no genetic variation. This presents a challenge when identifying and investigating symptoms of motor dysfunction, which can vary considerably in the human population. In order to seek variation in motor dysfunction, we have explored the use of a recombinant inbred family of mice known as the BXD panel, which combines well-characterized motor deficits within the BXD genetic reference panel. The BXD family, derived from the mating between C57BL/6J and DBA/2J parental strains, is the largest and best characterized mouse reference population, composed of ~150 lines (Ashbrook et al., 2021). The BXD strains have been very widely used to study the genetic basis of complex neurobehavioral traits such as anxiety, fear, and social behaviour (Philip et al., 2010). A major advantage of BXDs is that high-density genotype and phenotype data are publicly available (GeneNetwork, n.d.). The BXD strains of mice provide an opportunity to examine the genetic bases of variation in motor learning traits that are relevant to the phenotypes identified in individuals diagnosed with DCD.

A powerful approach to understanding the genetic basis of complex traits is through identification of Quantitative Trait Loci (QTLs). By combining molecular marker data of genetically related individuals with phenotypic trait values, genomic QTLs can be identified that contain genetic regulators of the trait.

QTL analysis can only be carried out in a population that differ genetically and phenotypically, such as the BXD RI lines of mice.

Since DCD is often detected in early childhood and persists into adolescence and adulthood, it is essential to follow the mice through the same timeline. Longitudinal research on disorders such as attention deficit hyperactivity disorder and autism spectrum disorder have demonstrated changes in clinical symptoms and underlying neurobiology across the lifespan, highlighting the importance of examining development longitudinally; but few studies have taken a developmental perspective when examining the genotype and phenotype of motor performance as it relates to neurodevelopmental disorders. Therefore, we conceptualized a battery of motor behaviour tasks across the mouse lifespan, as presented in a recent review by Gill et al. (2020). In our two-part study, we first presented the phenotypic and genotypic data for motor coordination in Gill et el. (submitted), and in this second part of the study we examined the natural variation in motor learning in BXD inbred strains of mice as it may relate to DCD. Specifically, the aims of this study were: (1) to determine whether the BXD recombinant strains of mice are a viable resource to assess variability in motor learning, (2) to utilize the QTL approach to evaluate the genome-to-phenome correlation in BXD strains of mice, and (3) to determine the translational value of the phenotypic and genetic findings for better understanding the human presentation of DCD.

## Materials & Methods

### Animals

All parents for each line were purchased from Jackson Laboratory (Bar Harbor, ME) and used to establish a colony at the Transgenic Animal Facility, Center for Molecular Medicine and Therapeutic, University of British Columbia. A total of 280 mice from C57BL/6J (B6), DBA/2J (DBA), and 12 BXD lines, ages postnatal (P)1-P120, were used in this study and housed in same-sex groups of 2-5 at a temperature of 22.5°C, with a 12:12h light-dark cycle (lights on at 6:00 AM PST). The number of mice tested for each task varied as mice were excluded if they were sick and/or injured. Further, the number of mice tested on the Complex Wheel and Skilled Reaching differed to ensure that the same mice that underwent dietary restrictions for skilled reaching task from each line were not tested on the complex wheel or horizontal rung task to avoid the effects of weight loss and motivation on motor learning. Therefore, the same mice that were administered the skilled reaching task did not take part in complex learning and vice versa. The total number of mice tested for each task are outlined in Table 1.

**Table 1:** Number of mice tested for each motor learning task.

| ID | Strain | Total number of mice in accelerating rotarod | Total number of mice in horizontal ladder rung | Total number of mice in complex wheel | Total number of mice in skilled reaching |
| --- | --- | --- | --- | --- | --- |
| 1 | C57BL/6J | 18 | 5 | 7 | 4 |
| 2 | DBA/2J | 18 | 7 | 9 | 4 |
| 3 | BXD1 | 18 | 17 | 8 | 4 |
| 4 | BXD15 | 18 | 16 | 11 | 4 |
| 5 | BXD27 | 18 | 18 | 9 | 4 |
| 6 | BXD28 | 18 | 11 | 11 | 4 |
| 7 | BXD32 | 24 | 21 | 11 | 4 |
| 8 | BXD40 | 24 | 18 | 12 | 4 |
| 9 | BXD45 | 16 | 16 | 6 | 4 |
| 10 | BXD65a | 27 | 27 | 14 | 4 |
| 11 | BXD69 | 25 | 17 | 12 | 4 |
| 12 | BXD75 | 22 | 21 | 4 | 4 |
| 13 | BXD81 | 18 | 16 | 8 | 4 |
| 14 | BXD86 | 23 | 23 | 7 | 4 |

Food and water were freely available, and bi-weekly cage changes took place after testing was complete for the week. All testing took place during the morning hours (light phase) between 8:00-12:00 PST by the authors KG and JR. All animal experiments were approved by the Animal Care Committee (ACC) of the University of British Columbia.

### Selection of BXD lines

The BXD family of recombinant inbred (RI) strains of mice were used for this study. The BXD strains are derived from crosses between two parental lines, C57BL/6J (B6) and DBA/2J (DBA), and these progeny were then inbred for at least 20 generations.. Twelve BXD RI lines (BXD1, BXD15, BXD27, BXD28, BXD32, BXD40, BXD45, BXD65a, BXD69, BXD75, BXD81, BXD86) were selected based on hypothesis driven data, which included selecting lines based on behaviour and structural brain differences similar to what is seen in children with DCD. As seen in Table 2, BXD1, BXD15, BXD27, BXD28, and BXD40 were selected due to their variation in cerebellar volume (Badea et al., 2009). Other strains were selected based on behavioural performance on motor coordination (BXD32, BXD45, BXD651, BXD69, and BXD75) and learning tasks (BXD81 and BXD86) that ranged from poor to very good, in relation to human DCD phenotypes (Philip et al., 2010) (Table 2). Common assays, such as rotarod, open field, and the dowel test, were used to select strains that demonstrated core DCD-like phenotypes ranging from very good to poor for balance (latency to fall on rotarod, dowel test), motor coordination (spontaneous locomotion in open field), and cerebellar involvement (cerebellar volume using MRI). In this manner, BXD69, BXD75, BXD81, and BXD86 were selected for their performance variability ranging from poor to very good on the rotarod (Brigman et al., 2008). The BXD32, BXD45, and BXD65a lines were selected either due to observed variation in balance abilities on the dowel test and/or spontaneous locomotion in the open field (Brigman et al., 2008; Philip et al., 2010).

**Table 2:** Selection of BXD lines based on phenotypes.

**Table Error! No text of specified style in document. Selection of BXD lines based on phenotypes**
| GeneNetwork ID | Phenotype | Lines from Jackson Laboratory |
| --- | --- | --- |
| 10921 | Cerebellum volume | BXD1(↑); BXD15(↔); BXD27(↓) |
| 10921; 11819 | Cerebellum volume; Dowel test | BXD28(↑) |
| 11819 | Dowel test | BXD32(↔) |
| 10921 | Cerebellum volume | BXD40(↑) |
| 11014 | Open field behaviour | BXD45(↑); BXD65a(↔) |
| 11004 | Rotarod performance | BXD69(↓); BXD75(↔) |
| 11014; 11005 | Open field behaviour; Improvement on rotarod | BXD81(↓) |
| 11004; 11819 | Rotarod performance; Dowel test | BXD86(↓) |

The selection of strains that demonstrated core phenotypes ranges between various levels (↑- high; ↔- medium; ↓- low)

### Procedures: Behavioural Testing

Detailed information on the study timeline is available in Figure 1 and rationale and procedural details for the behavioural tasks are outlined in Gill et al. (2020). This paper focuses on the first third and last phases of the study, as outlined in Figure 1. In the adult phase of testing, functional motor tasks were used to study motor learning in mice. In this phase, accelerated rotarod, horizontal rung walking, complex wheel, and skilled reaching tasks were carried out between P90-P120. The order of testing for all motor tasks was randomized for each litter on each testing session to limit the test order interaction and the effect of fatigue on performance. Further, we intentionally studied age-matched male-female pairs from different litters in each of the 14 strains. This ensured that the within-strain variance was not a litter effect only.

**Figure 1.**
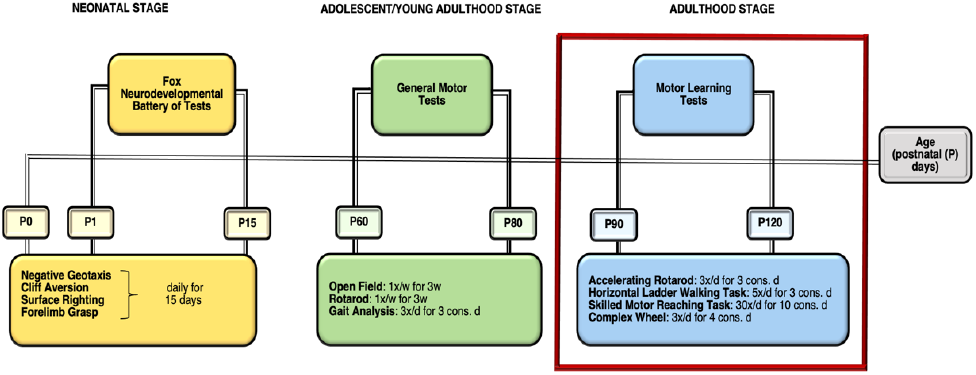
Workflow of the behavioral testing in Postnatal Day (P) P1-P120. All three phases of testing are proposed based on the DCD-like behavior. [d-days, w-week, cons.-consecutive]

### Statistical Analysis

#### Behavioural Data Analysis

Statistical analysis was carried out using IBM SPSS Statistics Version 25.0. Data were analyzed separately for each test by factorial ANOVA using line as an independent variable. Post hoc comparisons were performed when appropriate using the Bonferroni method. Due to potential litter effects, data used in analyses were the means of the line for each respective variable.

An average of all trials per day for each task was taken for each of the testing days, for each task. The effect of the line was explored using a two-way ANOVA with body weight and sex as covariates. To examine motor performance improvements and/or motor learning, a difference score was calculated (mean of last day of testing - mean of first day of testing) when appropriate. All measures are reported as the mean +/-S.E.M. and the level of significance reported for all comparisons is p < 0.05.

### QTL mapping and candidate gene analysis

All behavioral trait data were uploaded in GeneNetwork (www.genenetwork.org), an open access online database which contains BXD strain genomic and phenotypic information. These analyses classified the BXD strains in line with their genotypes using distinct chromosomal markers and compared them individually with the phenotypic variables. The likelihood ratio statistic (LRS) was computed to assess the strength of genotype-phenotype associations and identify QTLs that modulate a phenotypic trait. A test of 2000 permutations was performed to evaluate the statistical significance of associations. The bootstrap test was implemented to identify QTL location. A significant QTL is determined by the LRS (likelihood ratio statistics) value that corresponds to a genome-wide p-value of less than or equal to 0.05, whereas a suggestive QTL represents the LRS value that corresponds to a genome-wide p-value of less than or equal to 0.63. Confidence intervals around the significant LRS score peaks were calculated using 1.5 logarithm of the odds score.

The QTL maps were generated using GeneNetwork. The expression, functional and phenotypic information for each of the genes located within the significant QTL (Bello, Smith, & Eppig, 2015; Eppig, et al., 2015) was surveyed using Allen Brain Atlas, Mouse Genome Informatics (MGI) and PubMed. Literature and database searches were conducted to determine if each a gene under the significant QTL peak had a previously reported role in motor skills and/or motor learning. GeneNetwork variant browser was used to identify nonsynonymous polymorphisms. We also identified previously published, overlapping loci relating to abnormal phenotypes within our mapped QTL interval using the MGI resource (http://www.informatics.jax.org/allele).

## Results

### Motor Learning: Variability in BXD lines

#### Accelerated Rotarod

We determined rotarod performance by measuring the latency to fall from the rotarod. A two-way ANOVA revealed a significant interaction between the BXD lines and day of testing [F (13, 280) = 2.936, p < 0.001]. To calculate motor performance improvements, the rotarod performance on day 3 was subtracted from the performance on day 1 (mean of day 3 - mean of day 1). There were main effects of BXD strain on motor learning [F (13, 280) = 2.936, p=<0.001]. As seen in Figure 2, BXD27, BXD28, BXD65a, BXD75, and BXD86, along with the parental line DBA, had the lowest latency to fall on the first day of testing and also some of the lowest motor learning rates over the 3 days of testing. Additionally, BXD40 and BXD45 performed close to or at the average mark for both day 1 and day 3 of testing; however, they both had relatively low rates of motor learning from day 1 to day 3. There was no correlation between adult body weight and motor learning and no sex differences were observed on rotarod performance (p > 0.05).

**Figure 2.**
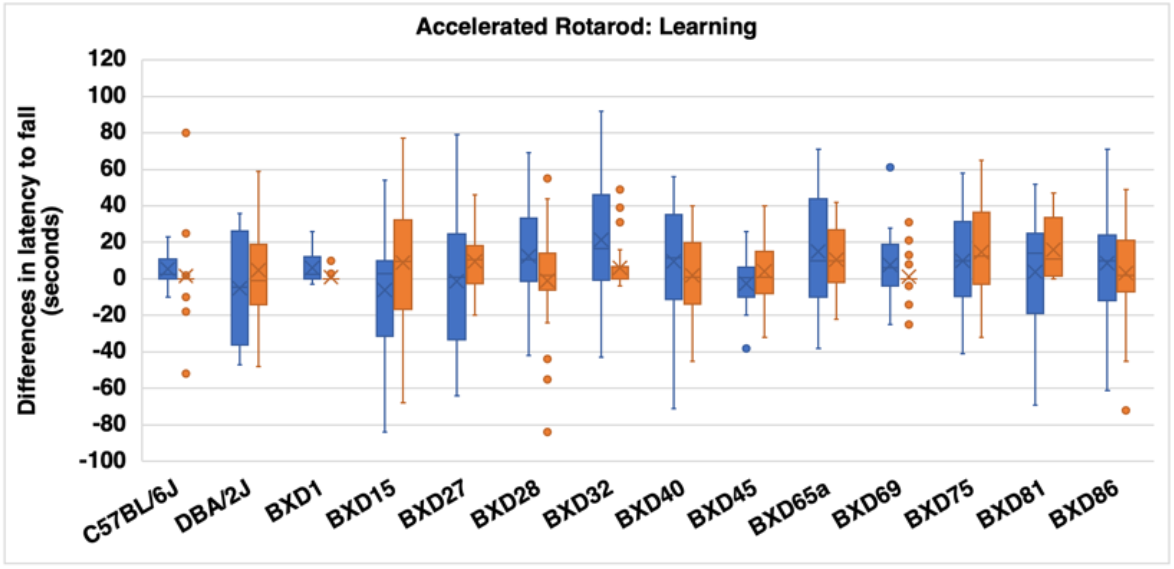
Graphs illustrating strain differences in accelerated rotarod parameter: learning. Blue represents day 1 strain performance in latency to fall, Orange represents day 3 strain performance in latency to fall. The x in each box depicts the mean and the bottom line of the box depicts the median.

### Horizontal Ladder Rung

The horizontal ladder rung walking task allowed for the measurement of generalizing motor learning from one day to the next. This task measured the rate of successful steps taken (number of correct steps taken/total number of steps taken); rate of correction (number of missteps self-corrected/total number of steps); and rate of missteps or errors (number of missteps/total number of steps) from baseline (day 1) to the last day of testing (day 3), indicating motor learning. Using a repeated measures ANOVA, there was significant interaction between test days and the BXD strain for the number of successful steps taken [F (13, 280) = 1.91, p < 0.05], rate of corrections made [F (13, 280) = 1.73, p < 0.05], and number of errors made [F (13, 280) = 4.22, p < 0.001]. There were main effects of the BXD line on number of successful steps taken [F (13, 280) = 4.89, p< 0.001], rate of corrections made [F (13, 280) = 5.61, p< 0.001], number of errors made [F (13, 280) = 9.27, p < 0.001]. Further, the fore-& hindlimb placement accuracy was measured, and it was found that for accuracy, limb placement correction, and error in fine and gross motor performance there was no statistically significant difference between [F (13, 219) = 5.028, p <.230] or within [F (101.64, 1712.2) = 1.347, p < 0.074] strain performance.

Although all strains performed relatively well on the horizonal rung walking test, as illustrated in Figure 3, BXD32, BXD40, BXD69, and BXD75 performed below average on the first day of testing. When assessing motor learning, BXD1, BXD28, and BXD65a were amongst the poorer learners. Although BXD32, BXD40, and BXD75 performed poorly on initial testing day, they were able to show the greatest improvements on the last day of testing. As in other measures, there was no effect of sex or weight on any of the above-mentioned measures (p > 0.05).

**Figure 3.**
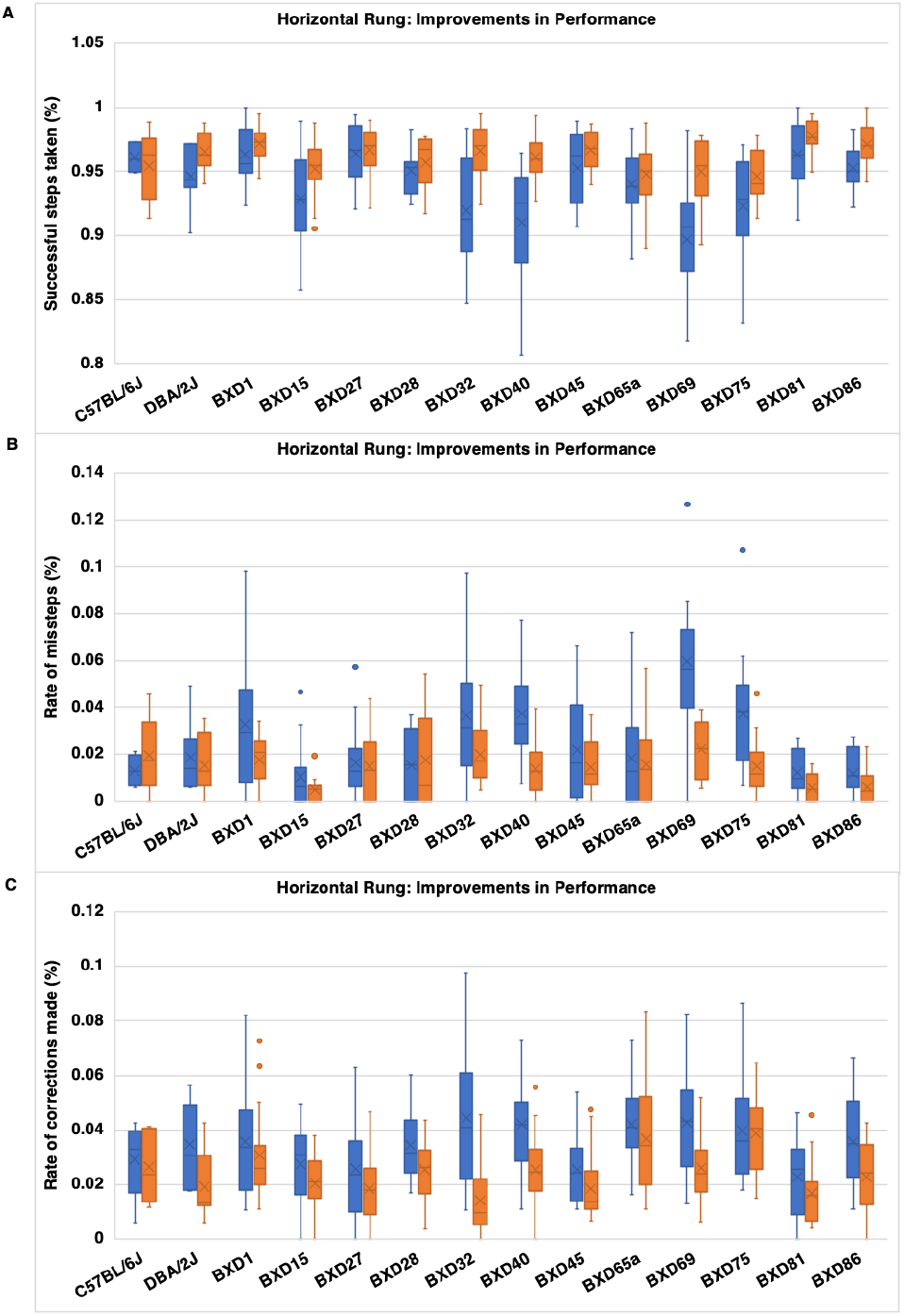
Graphs illustrating strain differences in horizontal ladder rung parameters: (A) successful steps take; (B) rate of missteps; (C) rate of corrections made. Blue represents day 1 strain performance in horizontal ladder rung, Orange represents day 3 strain performance in horizontal ladder rung. The x in the box depicts the mean and the bottom line of the box depicts the median.

### Complex Wheel

The complex wheel task assessed for motor learning with changing external task demands transference of motor learning from one day to the next. Using a two-way repeated measures ANOVA, the results revealed that there was an interaction between testing days and BXD strain for latency to fall from the wheel [F (13, 115) = 1.73, p < 0.005], as well as a main effect of BXD strain on the latency to fall [F (13, 115) = 16. 73, p < 0.001] (Figure 4A). A post-hoc analysis revealed that B6 was the only strain that significantly differed from all other strains, with no other strain showing a significant difference (p>0.05). Figure 4B shows that B6 outperformed all strains both during initial and final day of testing. Specifically, it’s seen that all lines, excluding B6 and BXD40 performed well below the average on the first day of testing. As seen in Figure 4, all lines, excluding BXD75, show improvements from first to last day of testing; however, not all lines show linear improvements. For example, lines such as BXD27 and BXD28 show an upwards trajectory from one day to the next in latency to fall (linear improvements); whereas lines such as BXD15, BXD40, an BXD86 vary from one day to the next.

**Figure 4.**
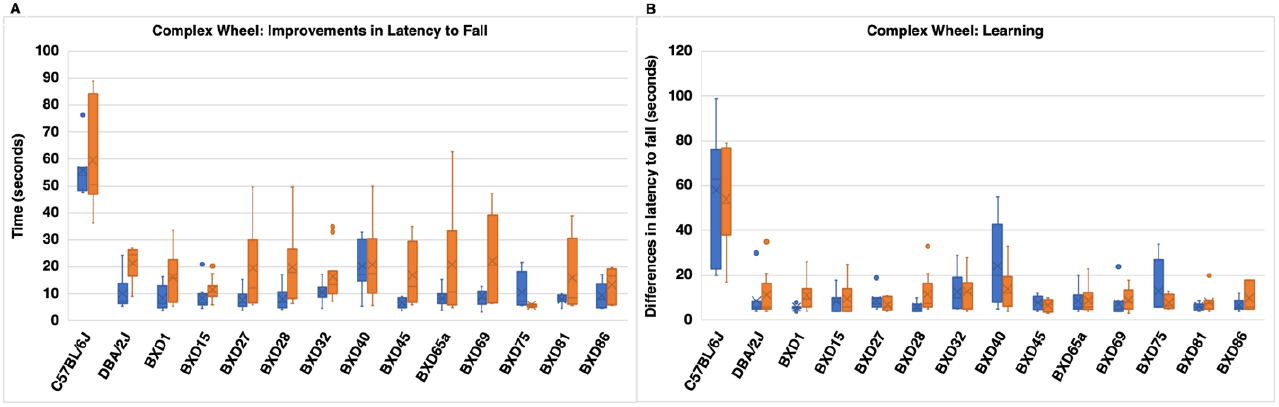
Graphs illustrating strain differences in complex wheel parameters: (A) latency to fall, Blue represents day 1 strain performance in complex wheel, Orange represents day 4 strain performance in complex wheel; (B) learning, Blue represents first day of trial 1 performance in complex wheel, Orange represents first day of trial 3 performance in complex wheel.. The x in the box depicts the mean and the bottom line of the box depicts the median.

When assessing the inter-trial learning within the same day of testing, statistically significant differences were observed [(F (13,115) = 2.214, p<.013)]. Specifically, BXD1 displayed short-term (i.e., intertrial) improvement within the same day of testing whereas BXD75, BXD40 and BXD86 displayed the least improvements within the same day. Further, motor performance improvement across the four-day testing period was calculated by subtracting final testing day (day 4) from the baseline (day 1). Using a one-way ANOVA, there was no significant difference in motor improvements between lines [F (13, 115) = 1.18, p = 0.304]. There was no effect of sex or weight on any of the above-mentioned measures (p>0.05).

### Skilled Reaching Task

This task measured complex motor learning of fine motor skills. This task involved food restriction, and consequently weight loss that served to be motivation to grasp food pellets. As there was a significant difference between lines for percentage of total weight loss [F (1, 55) = 2.52, p = 0. 012], weight loss was used as a covariate in this analysis. There were no statistically significant differences between lines for learning rate [F (1, 55) = 1.69, p = 0. 101], average first-time success [F (1, 55) = 1.65, p = 0. 109], or average total success [F (1, 55) = 1.56, p = 0. 135]. Figure 5 illustrates, BXD27, BXD28, BXD69, and BXD86 performed well below average for both first time success and total success (ability to grasp pellets regardless of the number of attempts). BXD40, BXD45, and BXD81 outperformed all other lines for first time success (i.e., grasping the pellet on the first attempt). Although not significant, it was found that all but one of the lines had higher total success when compared to first time success; however, BXD40 was the opposite, with higher first-time success.

**Figure 5.**
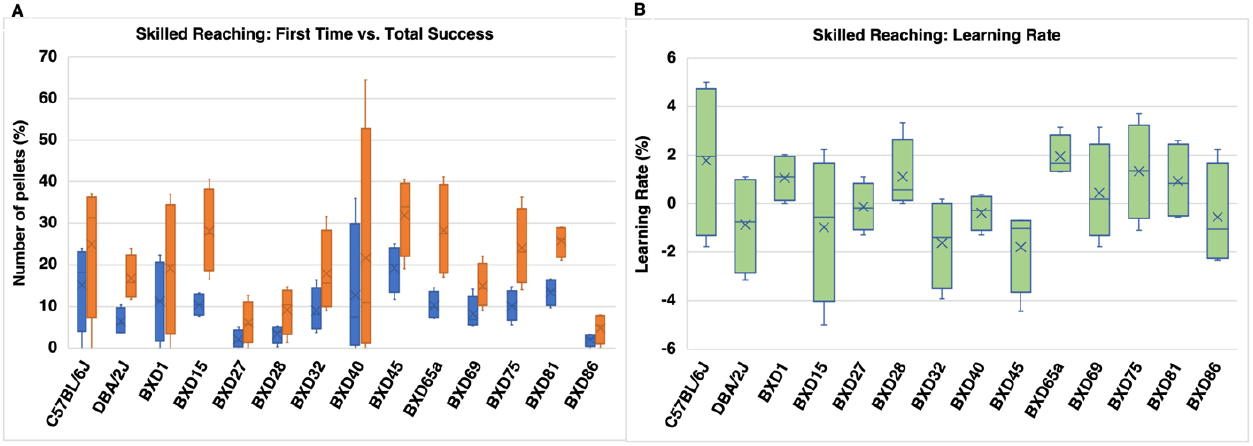
Graphs illustrating strain differences in skilled reaching parameters: (A) first time vs. total success; (B) learning rate.

### Heritability

For each of 11 phenotypes, heritability was calculated using an ANOVA with strain, sex, and weight as predictors. Heritability was calculated as the proportion of total variation explained by strain.

Phenotypes with high heritability have a large effect of genometype on the phenotypic variation. We can see that approximately 45% of the variation in weight is explained by strain. It is easier to identify QTL for traits with high heritability, since power to detect QTL is dependent upon heritability, locus effect size, number of genotypes examined, and number of biological replicates. We can see that even for phenotypes with low heritability (e.g. Horizontal Rung phenotypes) with sufficient numbers of individuals measures the strain effect becomes significant. Inversely, phenotypes where only a small number of animals were measured can still have high heritability.

### Underlying Genetics: QTL Analysis

#### Accelerated Rotarod

The QTL mapping for motor learning and balance phenotypes was performed using 12 BXD RI lines. The genome-wide QTL map for the difference in latency to fall parameter identified a single significant QTL to the proximal end of Chr 7 at 10.3 to 13.1 Mb with an LRS score of 22.21 [Trait ID 24722] (Figure 6).

**Figure 6.**
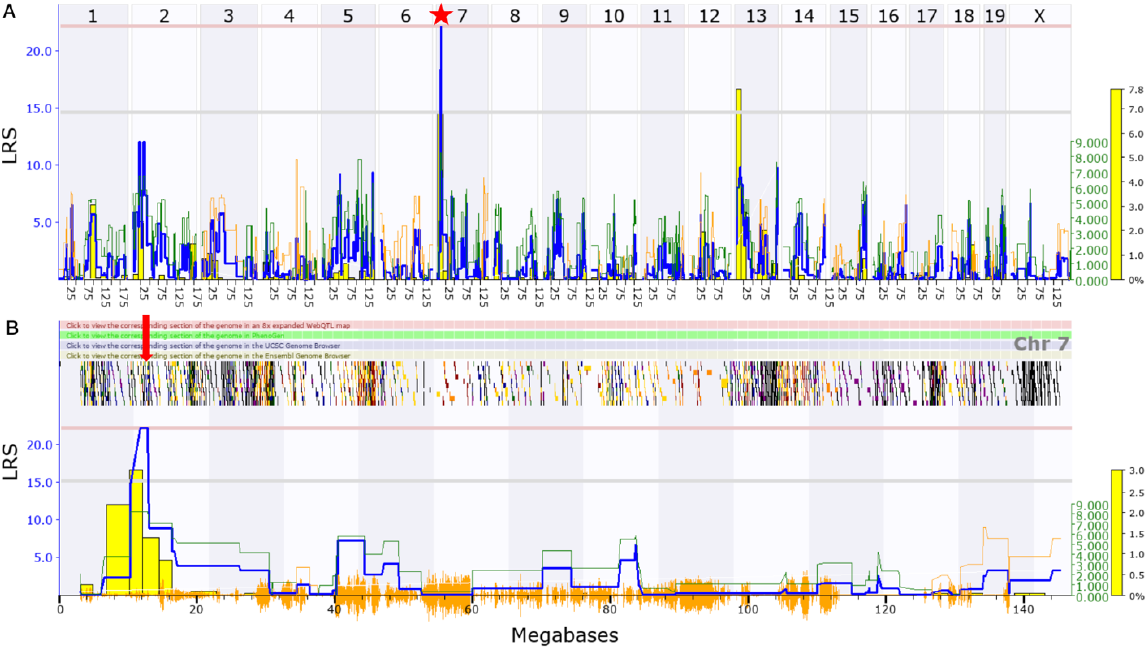
Genome-wide linkage map of learning (top to bottom) from accelerated rotarod to determine balance and motor learning. The overall blue trace shows the LRS. (A) Genome-wide QTL map showing a significant QTL on chromosome 7 for learning. (B) Interval QTL map of chromosome using three consecutive test day performance with bootstrap analysis. The lower gray horizontal line represents suggestive LRS genome-wide threshold at p ≤ 0.63. The upper pink horizontal line represents significant LRS genome-wide threshold at p ≤ 0.05. The bottom orange marks indicate SNP density. [asterisk (*) indicates significant QTL; down arrow (↓) indicates a higher resolution look at the QTL interval].

### Horizontal Rung Walking Task

The genome-wide QTL map for rate of missteps in fore- and hind-limb performance showed a significant QTL [Trait ID_24724] (Figure 7). This QTL was located on Chr 8 and spans a fairly small region from 66.9 to 72.55 Mb with an LRS value of 15.2. Additionally, there was the presence of suggestive QTLs on Chr 9 for missteps; and Chr 18, Chr 19, Chr 6 and Chr 8 for correction [Trait ID_24725] and successful steps taken [Trait ID_24723] (Figure S1).

**Figure 7.**
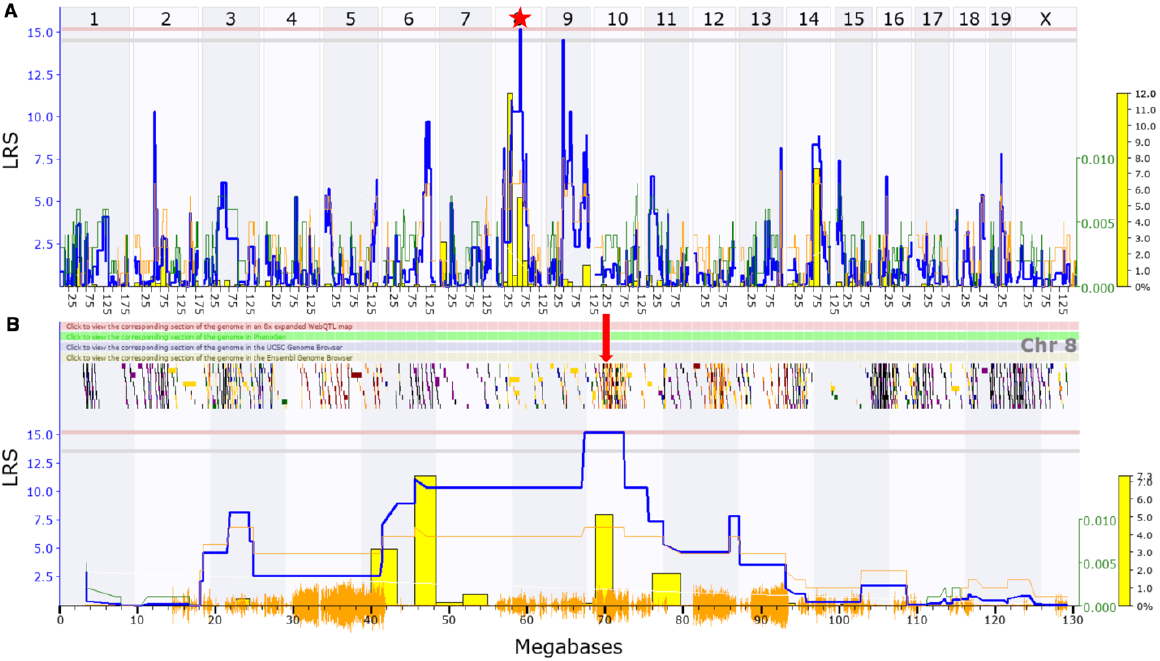
Genome-wide linkage map of rate of missteps (top to bottom) from horizontal ladder rung to determine number of errors in limb placement. The overall blue trace shows the LRS. (A) Genomewide QTL map showing a significant QTL on chromosome 8 for rate of missteps. (B) Interval QTL map of chromosome using three consecutive test day performance with bootstrap analysis. The lower gray horizontal line represents suggestive LRS genome-wide threshold at p ≤ 0.63. The upper pink horizontal line represents significant LRS genome-wide threshold at p ≤ 0.05. The bottom orange marks indicate SNP density. [asterisk (*) indicates significant QTL; down arrow (↓) indicates significant QTL interval with genes]

### Complex Wheel

The genome-wide QTL mapping for latency to fall did not result in any significant QTLs; but did identify suggestive QTLs on Chr 14, Chr 4, Chr 6 and Chr 15 [Trait ID 24729] (Figure S2).

### Skilled Reaching Task

The QTL mapping for skilled motor movements was performed separately for each parameter - first attempt success, total success and learning rate. The genome-wide QTL map identified a significant QTL for first attempt success (Figure 8). The significant QTL was located at the proximal end of Chr 15 [Trait ID_24726] and spans about 17 to 18.9 Mb with an LRS score of 20.89. In addition, a suggestive QTL for total success [Trait ID_24727] were found at Chr 15 (Figure S3).

**Figure 8.**
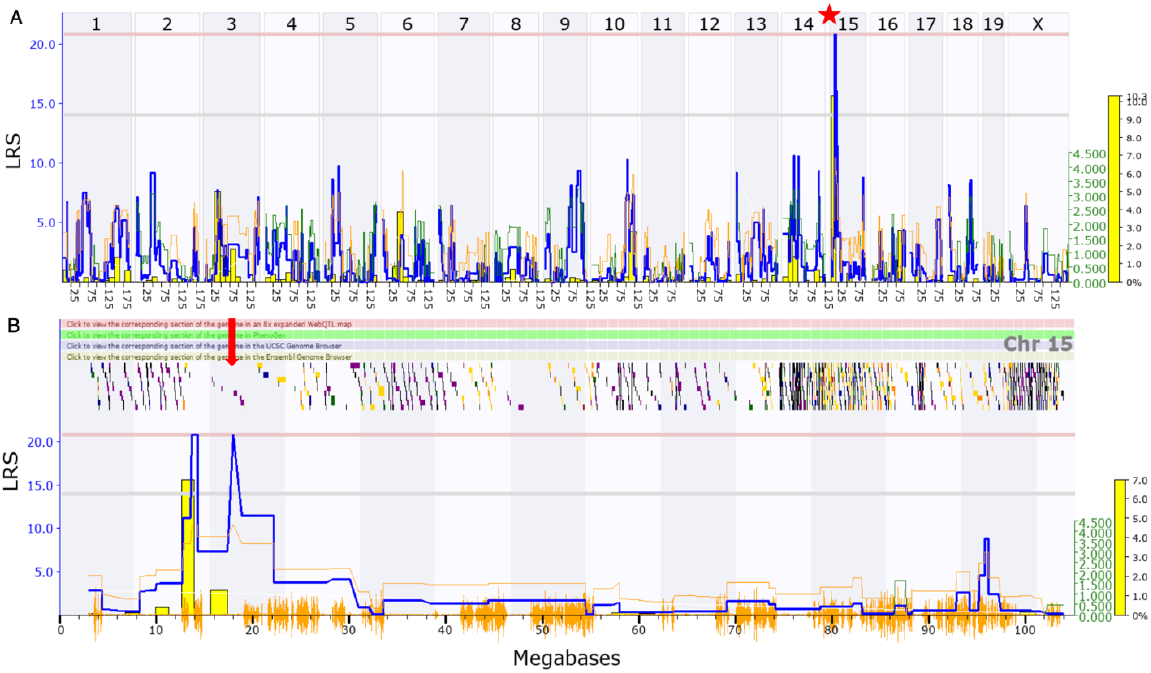
Genome-wide linkage map of first-time success (top to bottom) from skilled reaching to determine skilled motor movements. The overall blue trace shows the LRS. (A) Genome-wide QTL map showing a significant QTL on chromosome 15 for first time success. (B) Interval QTL map of chromosome using ten consecutive test performance with bootstrap analysis. The lower gray horizontal line represents suggestive LRS genome-wide threshold at p ≤ 0.63. The upper pink horizontal line represents significant LRS genome-wide threshold at p ≤ 0.05. The bottom orange marks indicate SNP density. [asterisk (*) indicates significant QTL; down arrow (↓) indicates a higher resolution look at the QTL interval].

### Candidate Genes for Significant QTLs

Following the successful identification of significant QTLs for three traits in the accelerated rotarod, horizontal ladder rung and skilled reaching tasks (Table 3), we focused on the best candidate genes underpinning each QTL. We find a total of 218 genes that mapped to the three significant QTL regions (Table 4). As discussed in the Materials and Methods section, we used a series of criteria to prioritize promising candidates: (A) the gene must be expressed in central nervous system (CNS) and/or skeletal muscle; (B) the gene must be associated with a motor function related to DCD-like behavior; and (C) presence of nonsynonymous SNPs in coding region of the gene. The expressed sequence tags and Riken clones within the gene list were not evaluated since they are currently non-annotated in terms of expression pattern and function. Of all the genes observed across three significant QTLs, 155 genes met criterion A (Table S1); 1 of these 155 genes met criterion A and B (Table 4); and no genes met all three criteria (Table S1). Tissue-specific expression of genes is crucial in understanding the role of a gene in a given disorder. Hence, we screened the 218 gene expression profile datasets to determine expression in brain and skeletal muscle tissues relative to other tissues using the Allen brain atlas, NCBI and MGI. Of the 218, we identified 155 genes that had relatively higher expression in CNS and skeletal muscles (Table S1). Secondly, the functional role of these 155 genes was explored using subject queries in PubMed and only identified *Rab3a* gene. *Rab3a*, however, did not have any non-synonymous polymorphisms (Table S1). Altogether, we have identified *Rab3a* as the only candidate gene that met at least two criteria across the three significant QTL regions.

**Table 3:** Heritability (h^2^) of phenotypes. Heritability was calculated as the proportion of total variance explained by strain. The significance (p-value) and number of individuals phenotyped (n) is shown for each phenotype.

|  |  |  |  |
| --- | --- | --- | --- |
| Phenotype | $h^2$ | p-value | n |
| Weight | 0.448852809 | < 0E-8 | 276 |
| Horizontal Rung - Rate of missteps | 0.205140716 | 5.29E-07 | 233 |
| Horizontal Rung - Rate of successful steps taken | 0.21604558 | 6.73E-07 | 233 |
| Accelerated Rotarod - Motor Performance Improvements | 0.111265078 | 0.002528841 | 286 |
| Horizontal Rung - Rate of correction | 0.124955341 | 0.006732658 | 233 |
| Complex Wheel - Latency to Fall | 0.14659015 | 0.16268035 | 129 |
| **Accelerated Rotarod - Learning** | 0.05540936 | 0.292335039 | 286 |
| **Skilled Reaching - Learning Rate** | 0.342733099 | 0.394718524 | 56 |
| **Skilled Reaching - First Time Success** | 0.408420222 | 0.414290731 | 56 |
| **Skilled Reaching - Total Success** | 0.377000568 | 0.421257552 | 56 |
| **Complex Wheel - Learning** | 0.07449641 | 0.914508276 | 129 |

**Table 4.**
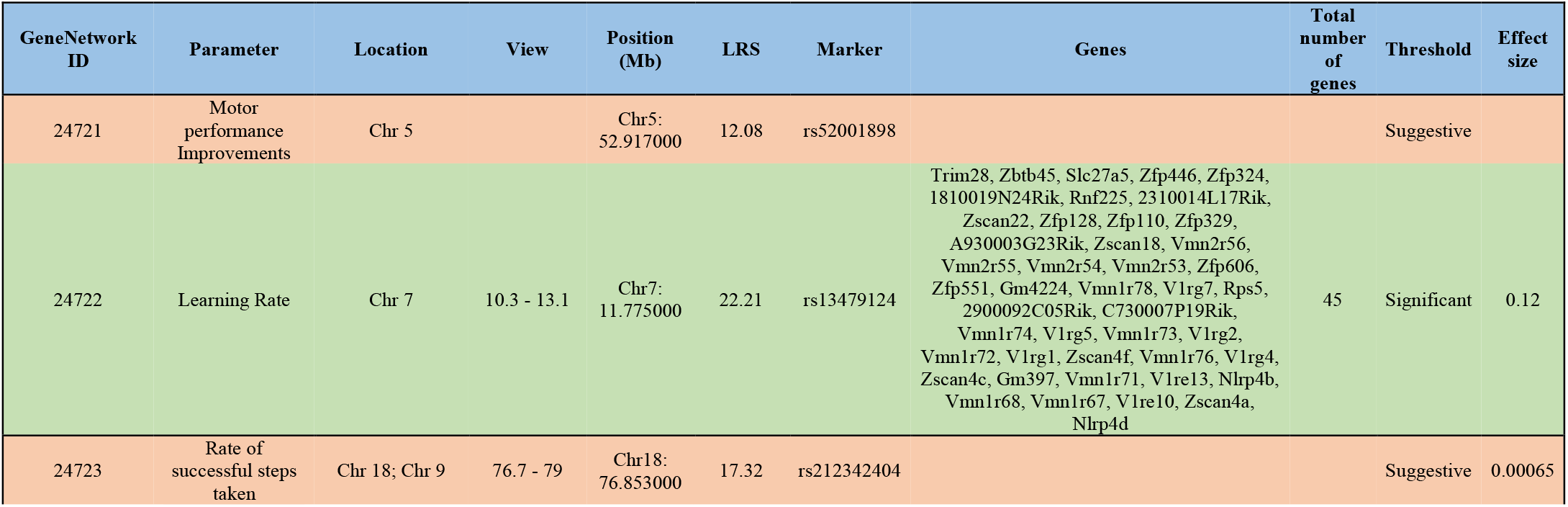
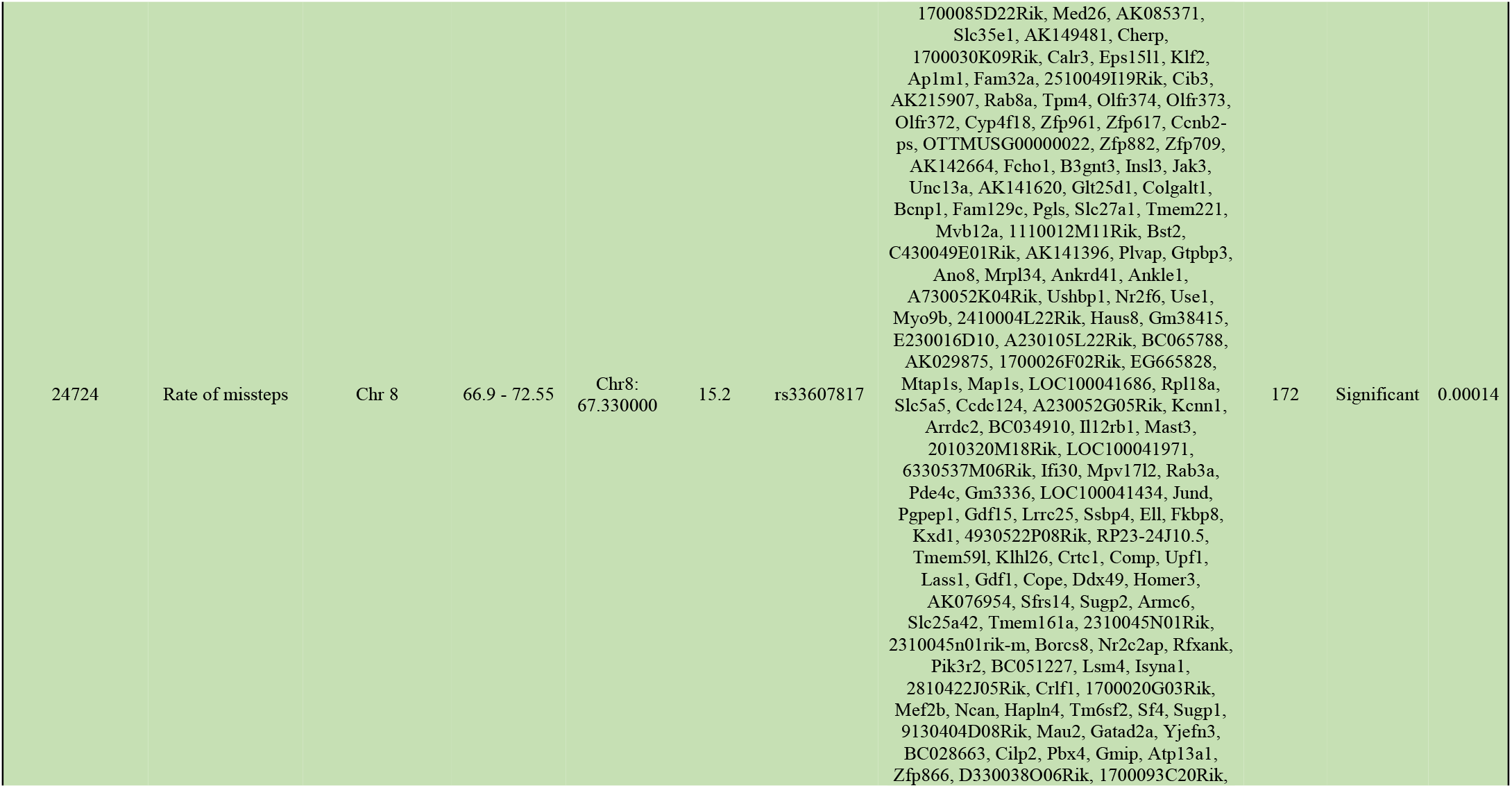
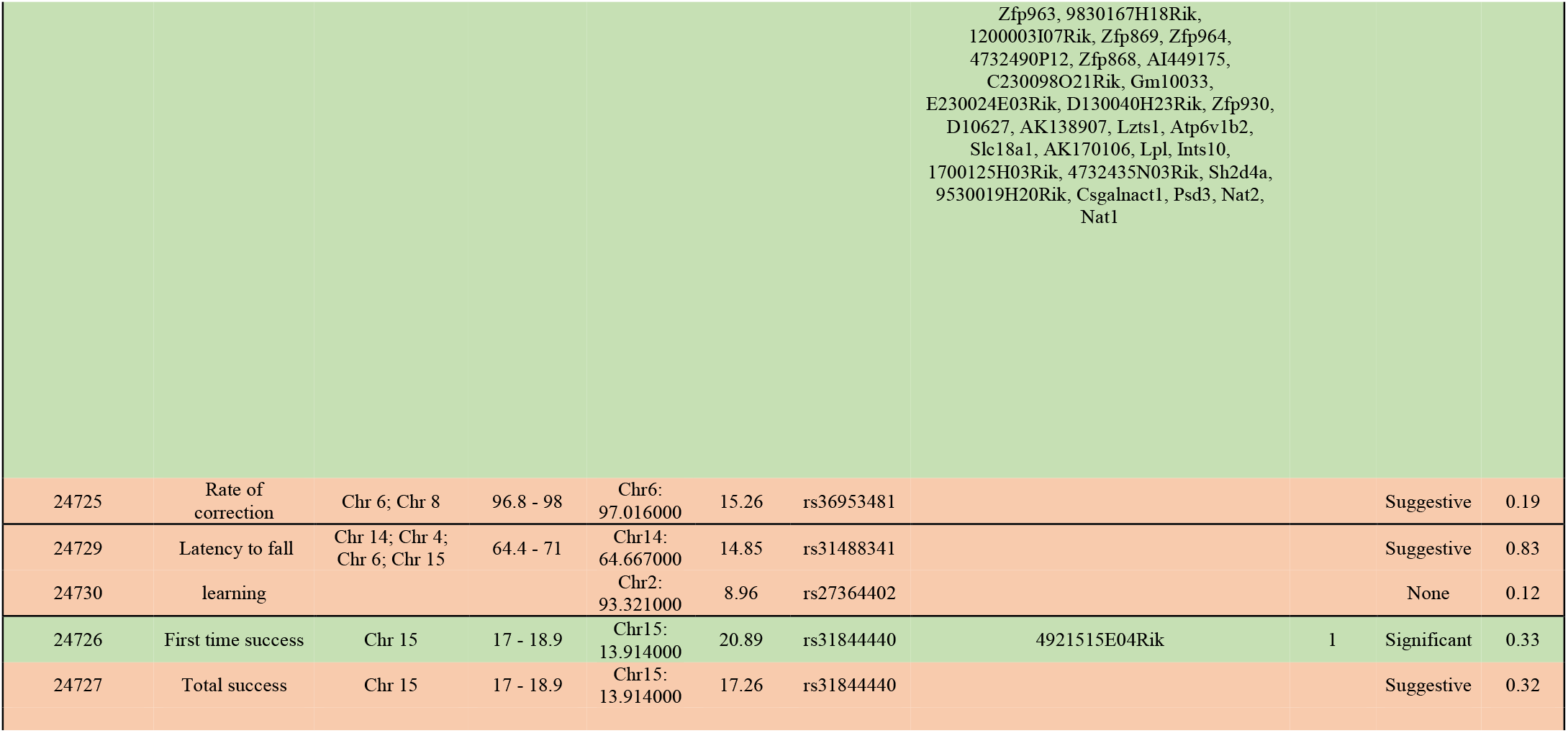
Summary of QTL analyses for motor learning measures. Each gene underlying the significant and suggestive QTL intervals is provided (green-significant; orange-suggestive)

## Discussion

The objective of this study was to examine the range of motor learning abilities in pre-selected BXD strains of mice to investigate whether we have access to phenotypes and genotypes that are relevant to DCD, and then to assess whether these phenotypic variations led to corresponding QTLs. The results of this study show that there is significant variability in motor learning parameters, as measured by accelerated rotarod, horizontal rung walking task, complex wheel, and skilled reaching task. The QTL analysis of skilled motor learning phenotypes identified four significant QTL peaks, one for each of the following parameters: motor improvement in accelerating rotarod, overall learning in fore- and hindlimb performance in horizontal ladder rung walking task, first attempt success in skilled motor reaching task, and inter-trial learning in complex wheel.

### Motor learning in BXD lines of mice

Results in this current study indicate there is a spectrum of motor learning in the pre-selected BXD strains of mice with a spectrum between high and low learning capabilities. The results indicate that certain lines struggle with motor coordination and motor learning, that is similar to DCD-like symptomology. It was found that BXD15, BXD27, BXD28, BXD75, and BXD86 presented with a varying motor profile, but all showed impairments with motor learning when compared to other strains. The motor learning impairments generally fit into the following categories: difficulties with (1) gross motor skills, (2) fine motor skills, or (3) both gross and fine motor skills.

Gross motor skills include large muscle movements involving the arms, legs, and the back that are functionally required for sitting, standings, walking, and running. As seen in Figures 2 and 4, BXD15 and BXD75 generally fit into the first category who primarily show gross motor impairments, measured by the accelerated rotarod and complex wheel tasks. Both strains struggled on tasks that required motor coordination of large muscle movements, dynamic balance, and locomotor skills. BXD15 and BXD75 also show poorer adjustment to changing external demands that require readjusting the body in space based on visual and kinesthetic information. Specifically, BXD75 had one of the lowest baseline performances and the least improvements on latency to fall on the accelerated rotarod, and the complex wheel. This lack of motor learning may be due to impairments in dynamic balance, and visual-spatial integration, as these tasks are highly demanding of balance and visual perceptual abilities (Silverman, 2010). Further, BXD15 also struggled with dynamic balance and motor learning that involved adjusting to external task demands, as evidenced by negative learning on the accelerated rotarod and one of the lowest learning abilities on the complex wheel. These tasks require changing task demands and require the animal to integrate spatial information with appropriate body adjustment (i.e., adjusting to the constant change in acceleration). Both lines, however, show close to or above average learning on motor tasks that require fine motor coordination and learning (i.e., horizontal rung walking task and skilled reaching).

Fine motor skills are defined as smaller actions, such as grasping an object between the thumb and a finger or using the lips and tongue to feed (Mon-Williams et al., 1999). In our model, the horizontal rung walking task and the skilled reaching task both require fine motor coordination and learning. We saw that BXD28 performed relatively well on the tasks requiring gross motor skills, such as latency to fall on the accelerated rotarod, which is similar to the finding of Philip et al. (2010). However, BXD28 had one of the lowest learning rates on both the horizontal rung and skilled reaching tasks (Figures 3 and 5). Further analysis of horizontal rung walking task indicated that all lines had some extent of self-correction ability; however, over time they had to self-correct less often and had lower error rate, consequently a higher success rate. However, BXD28 did not show the same pattern as there was not a decrease in error rate; however, there was a decrease in self-corrections made. In the case of BXD28, this did not translate to higher success rate, instead BXD28 had the lowest success rate over the three days of testing, indicative of poorer learning. Although it is not clear why BXD28 struggle with fine motor learning, it is possible that they persisted in using less efficient strategies and tend to repeat the same movements without making corrections to their performance. For the skilled reaching task, BXD28 had one of the lowest first-time success values, the ability to grasp the pellet from the skilled reaching box on the first attempt, and the lowest total success at grasping the pellet. This may be due to impairments in visual monitoring, and associated lack of terminal control of accuracy as they are not able to control their endpoint appropriately based on the visual feedback (Missiuna, 1992). Zou & William (1999) found that BXD28 are amongst the BXD strains with the lowest number of retinal ganglion cell numner. It is known that the retinal ganglion cells of the retina play a role in anticipating the location of smoothly moving objects such as, the pellet in the case of our study, and retina can also signal violations in it’s own predictions (Schwartz et al., 2007). Based on this evidence, it can be understood that BXD28 may present difficulties with learning fine motor skills due to impairements in the visual and neural system.

In this current study we found two lines that show mixed impairments in both fine and gross motor tasks. Similar to Philip et al. (2010) findings, in our study BXD27 and BXD86 show poor performance on the accelerated rotarod; however, in this study we also investigated the ability to learn over three consecutive testing days and found that BXD27 and BXD86 had relatively poorer motor learning abilities despite repeated practice. This indicates that both strains struggle with learning gross motor skills. When assessing performance on fine motor skills, BXD27 and BXD86 mice were amongst the poorest learners on horizontal rung walking task (Figure 3). Similar to BXD28 mice, as seen in supplementary Figure 2, BXD27 mice decreased the number of self-corrections made over time; however, this did not impact the number of errors over time. BXD27 mice continued to make a similar number of errors on the final day of testing when compared to the initial testing day. Consequently, BXD27 presented with a lower success rate and decreased motor learning over the testing days. This may indicate that BXD27 persist in movement strategies that may not be effective, possibly due to the inability to update the internal model of movement that occurs in the cerebellum (Badea et al., 2008; Ceccarelli et al., 2015; Galante et al., 2009). To further support this hypothesis, there is evidence to show that BXD27 have the lowest cerebellum volume [GeneNetwork Trait ID_1092] and retinal ganglion cell number [GeneNetwork Trait ID_10649; 10650] when compared to other BXD lines of mice (Badea et al., 2009; Schwartz et al., 2007). As stated initially, the cerebellum is believed to be the main neural correlate motor impairments (Zwicker et al., 2009). The cerebellum and the visual system, specifically the retina play a role in updating the internal model of movement in order to facilitate motor coordination, motor learning, postural control and cognition (Schwartz, 2007; Zwicker et al., 2009). Animal studies have suggested a causal relationship between motor skills acquisition and cerebellar volume (Marchiori, 1987; Steadman et al., 2014; Zwicker et al., 2010). Additionally, Figure 5 indicates that BXD27 and BXD86 had the lowest average first time success and the lowest average total success when grasping the pellet. This indicates that both BXD27 and BXD86 were amongst the poor learners on the gross motor learning tasks (i.e., accelerated rotarod and complex wheel; Figures 2 and 4), and the fine motor learning tasks (i.e., horizontal rung and skilled reaching; Figures 3 and 5). Similar to individuals with DCD, it is possible that BXD27 and BXD86 plan, monitor, and evaluate their performance less often, leading to greater impairments in motor execution on a global level.

### Motor behavior QTLs and previous QTL analyses

The functional roles of significant QTL genes were assessed in relation to DCD-like behavior. Only *Rab3a* (Ras-related protein Rab-3A) emerged as a best candidate gene for the horizontal ladder rung task. This gene is found to be associated with brain and skeletal muscle development (http://www.informatics.jax.org). Behaviorally, Rab3a disruption in the mouse has been shown to alter several behaviors, including circadian activity and sleep homeostasis (Kapfhamer et al., 2002), reversal learning and exploration (D’Adamo et al., 2004), regulation of emotion (Yang et al., 2006) and memory precision (Ruediger et al., 2011). These findings may provide a starting point for further studies of the motor skills deficits underlying impaired motor learning and the variations in motor phenotypes found in BXD lines of mice. It has been shown that DCD has an underlying genetic component due to its high heritability (~70%) (. Lichtenstein et al. (2010). Qian et al. (2013) found genetic variation in dopamine-related gene expression in relation to motor learning, suggesting that variation in dopamine receptors may play a role in DCD.

The BXD lines are a very rich and extensive genetic resource, and we have the opportunity to take advantage of this by using them to map traits, specifically motor traits, to lay the groundwork for the study of the genetic link to DCD-like phenotypes. In an effort to determine if there were any intersections or overlaps between our data and previously reported data within our mapped QTL interval of motor phenotypes, we used the Mouse Genome Informatics (MGI) resource. This resource provides a wealth of information regarding the mouse genome, including comprehensive gene maps, molecular and genetic markers, and phenotypic descriptions. We inputted all significant QTL locations [(Chr 7: 10.3 to 13.1 Mb); (Chr 3: 66.9 to72.55 Mb) and (Chr 15: 17 to 18.9 Mb)] individually for learning rate, number of correct limb placement that involved measuring balance, motor learning, rate of missteps and first-time success. The phenotype, alleles, and disease models query output generated various mutations such as targeted, transgenic, endonuclease-mediated and phenotype data. The chromosomal location of all three QTLs was also found associated with abnormal phenotypes that included skeletal muscle weight (Hernandez-Cordero et al., 2019); Body weight (Rocha JL, et al., 2004) and stress response (Thifault et al., 2008). Similarly, all suggestive QTL locations [(Chr 18: 76.7 to 79 Mb); (Chr 6: 96.8 to 98 Mb); (Chr 14: 64.4 to 71 Mb); (Chr 15: 17 to 18.9 Mb) and (Chr X: 131.4 and 135.8 Mb)] were keyed in for rate of successful steps taken, rate of correction, latency to fall and total success. The phenotype, alleles, and disease models query output generated abnormal phenotypes Body weight (Rocha JL, et al., 2004; Halliwill et al., 2016, Dragani et al., 1995); Postnatal body weight (Ishikawa et al., 2004) and locomotor activity (Downing et al., 2003). Based on the literature evidence, these abnormal phenotypes may be interrelated. For example, Cairney & Veldhuizen (2013) reported that physical inactivity among children with DCD increases their tendency toward obesity compared to typically developing children. The poor motor skills of children with DCD can create stress in physical activity settings (Zimmer, Dunn & Holt. 2020).

Altogether, these findings support that specific genes and/or gene networks with functional implication in motor behavior are associated with the development and function of neural circuits that are implicated in DCD. It is worth mentioning that a vast number of gene-phenotype relationships could be unreported (Meehan et al., 2017). In addition, many studies only focus on genes that are well-annotated and ignore many potentially important genes that are less well understood (Stoeger, Gerlach, Morimoto, & Nunes Amaral, 2018).

### DCD-like Phenotypes

Individuals with DCD present as a heterogeneous group of individuals who all have delayed acquisition of motor skills but show a wide range of motor coordination difficulties, similar to what was observed in the above mentioned BXD lines of mice (Missiuna, 1992). Children may demonstrate competence in some motor skills while impairments are noticed in others. Williams et al. (2008) suggests that the presentation of motor coordination and learning deficits can vary in individuals with DCD due to factors such as the individual’s level of motor impairments and task complexity. As mentioned earlier, individuals with DCD struggle with either fine motor skills, gross motor skills, or both. Drawing a parallel to BXD line of mice, BXD15 and BXD75 primarily presented with gross motor learning impairments, BXD28 with fine motor learning, and BXD27 and BXD86 demonstrate impairments in both gross and fine motor learning skills. In children, the heterogeneity in motor skills is usually evidenced on the leading assessment tool of DCD, Movement Assessment Battery for Children-2 (MABC-2) (Henderson et al., 2007; Blank et al., 2019). Some children are seen to struggle with gross motor skills, such as aiming and catching a ball and/or balance, while others have difficulties with final motor skills, including manual dexterity tasks. There are also a large percentage of children who show impairments in both gross and fine motor skills of the MABC-2. Similarly, this study used various tasks that require gross motor and fine motor skills to assess motor coordination and motor learning. The results are similar in our animal model as children with DCD in which there is a heterogenous profile of motor learning.

## Conclusion

Taken together, the evidence from this study suggests that certain lines struggle with motor coordination and motor learning. The BXD15, BXD27, BXD28, BXD75, and BXD86 variants presented with a varying motor profile, but all struggled with motor impairments compared to other lines. Five lines - BXD15, BXD27, BXD28, BXD75, and BXD86 - exhibited the most DCD-like phenotype from infancy to early adulthood, when compared to other BXD lines of interest. BXD15 and BXD75 struggled primarily with gross motor skills, BXD28 primarily had difficulties with fine motor skills, and BXD27 and BXD28 lines struggled with both fine and gross motor skills. In summary, the motor difficulties experienced by these lines need to be further analyzed to investigate the resemblance to DCD-like phenotypes. Additionally, it is suggested that future work expland the study to include more lines of BXD mice, which would provide more confidence in the QTL results. Future work will investigate neuroanatomy and phenotype-genotype linkage to investigate the neurobiological and genetic link for DCD.

## Supplementary Figures

**Figure S1.**
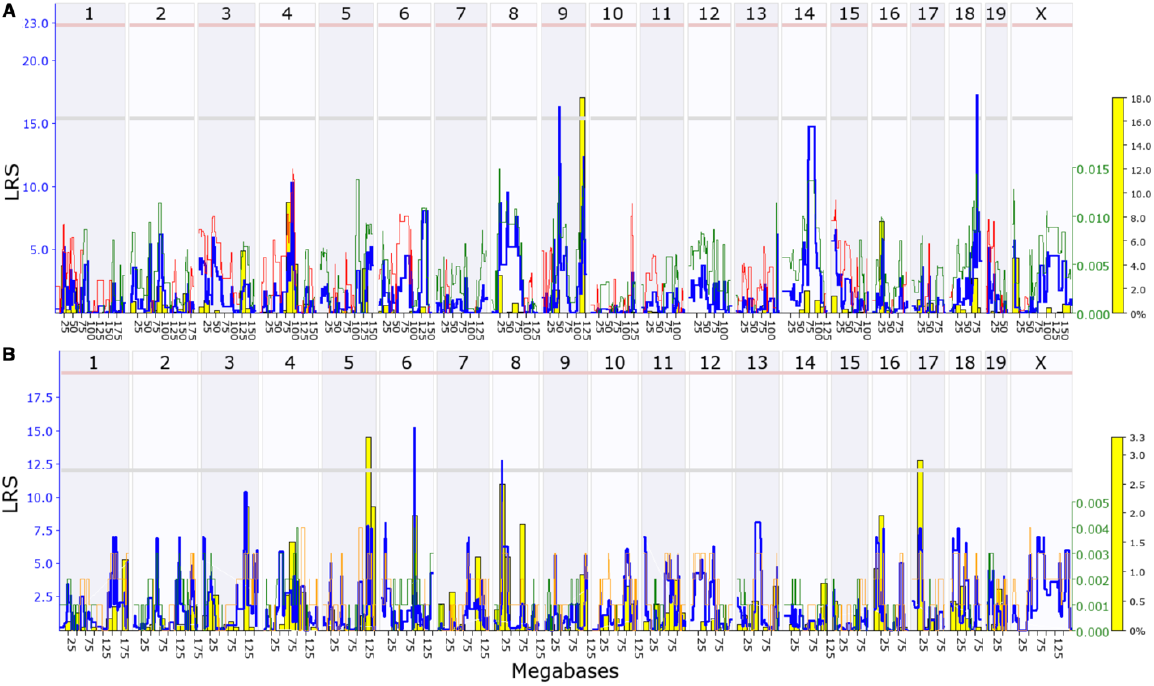
Genome-wide linkage map of (A) rate of successful steps taken and (B) rate of correction (top to bottom) from horizontal ladder rung to determine fore-& hindlimb placement accuracy and number of correct limb placement. The overall blue trace shows the LRS. The genome-wide QTL map showing suggestive QTLs on Chromosome 18, Chromosome 9 for rate of successful steps taken & Chromosome 6, Chromosome 8 for rate of correction. The lower gray horizontal line represents suggestive LRS genome-wide threshold at p ≤ 0.63. The upper pink horizontal line represents significant LRS genome-wide threshold at p ≤ 0.05. The bottom orange marks indicate SNP density.

**Figure S2.**
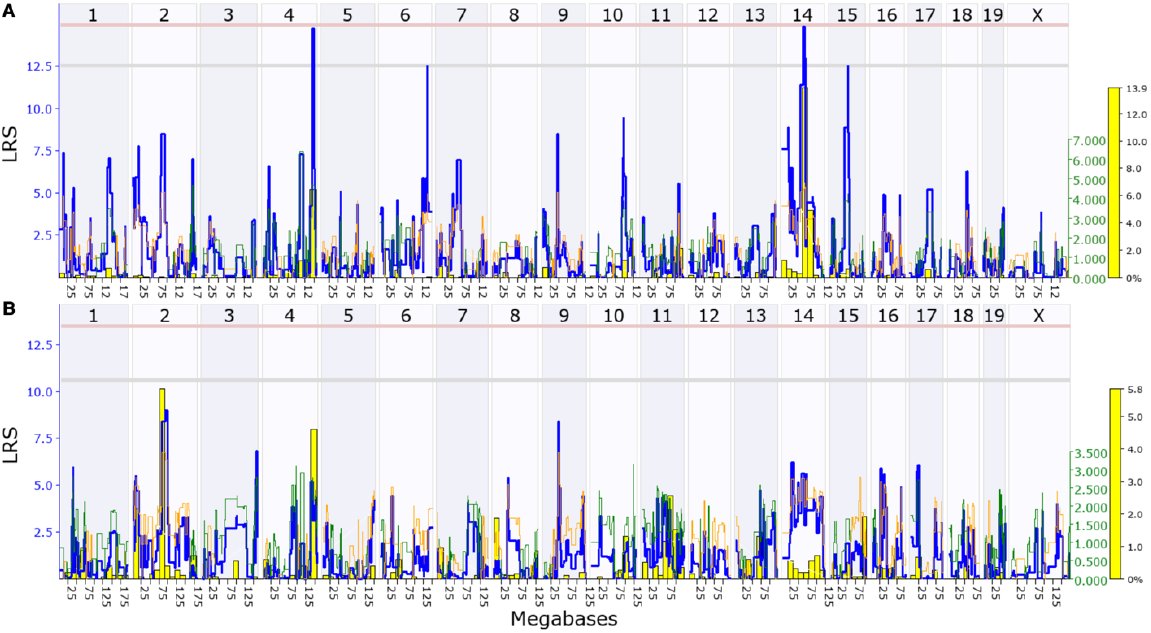
Genome-wide linkage map of (A) latency to fall and (B) learning (top to bottom) from complex wheel to determine noval motor learning. The overall blue trace shows the LRS. The genomewide QTL map showing suggestive QTLs on Chromosome 14 for latency to fall & no threshold for learning rate. The lower gray horizontal line represents suggestive LRS genome-wide threshold at p ≤ 0.63. The upper pink horizontal line represents significant LRS genome-wide threshold at p ≤ 0.05. The bottom orange marks indicate SNP density.

**Figure S3.**
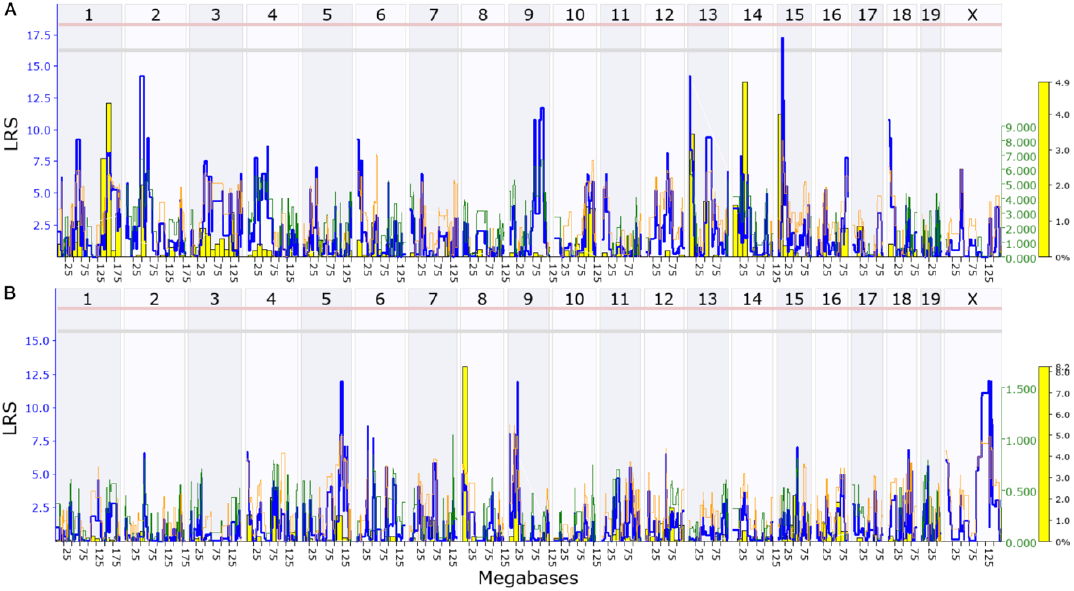
Genome-wide linkage map of (A) total success and (B) learning rate (top to bottom) from skilled reaching to determine skilled motor movements. The overall blue trace shows the LRS. The genome-wide QTL map showing suggestive QTLs on Chromosome 15 for total success & no threshold for learning rate. The lower gray horizontal line represents suggestive LRS genome-wide threshold at p ≤ 0.63. The upper pink horizontal line represents significant LRS genome-wide threshold at p ≤ 0.05. The bottom orange marks indicate SNP density.

**Table S1.**
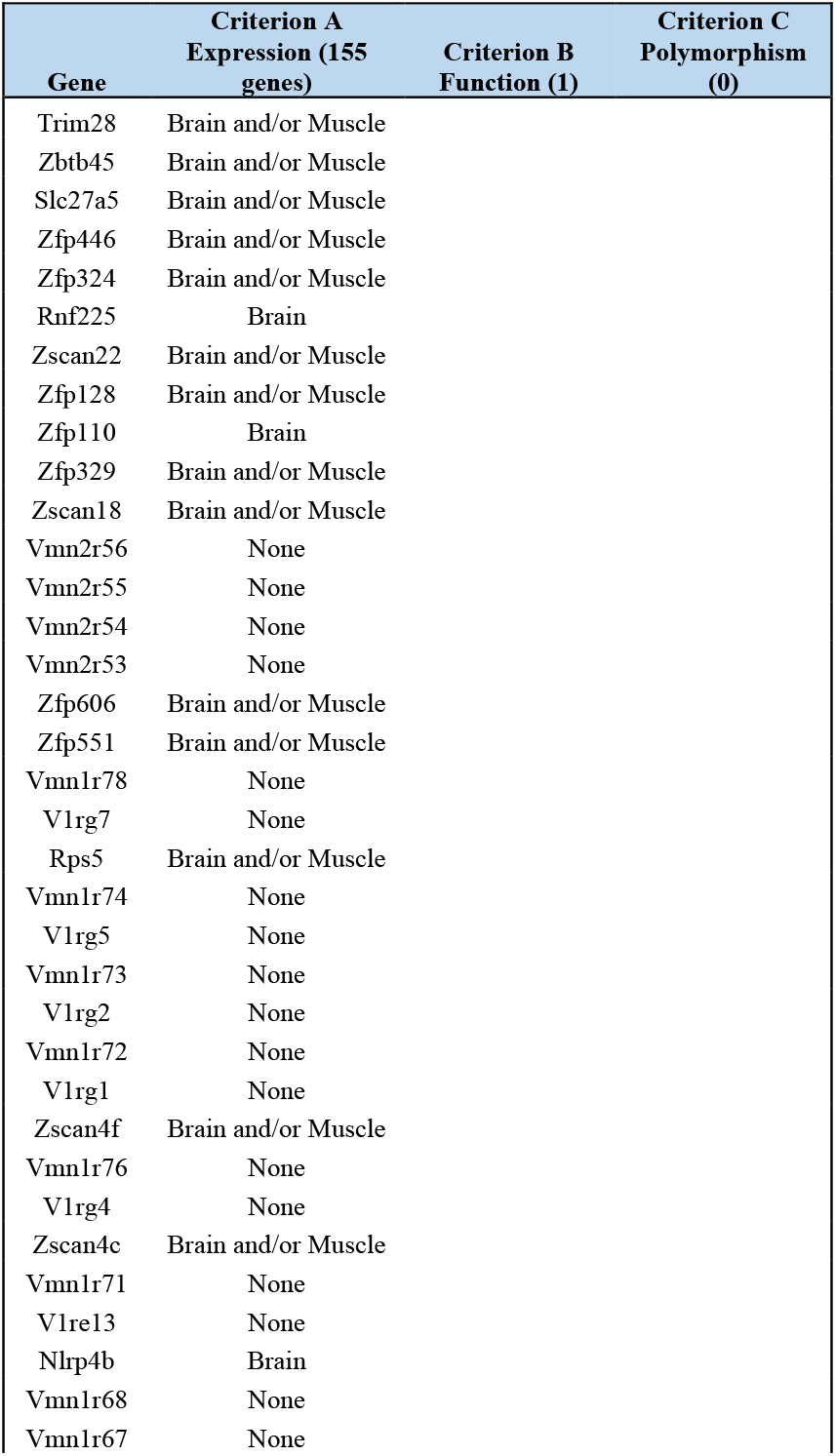
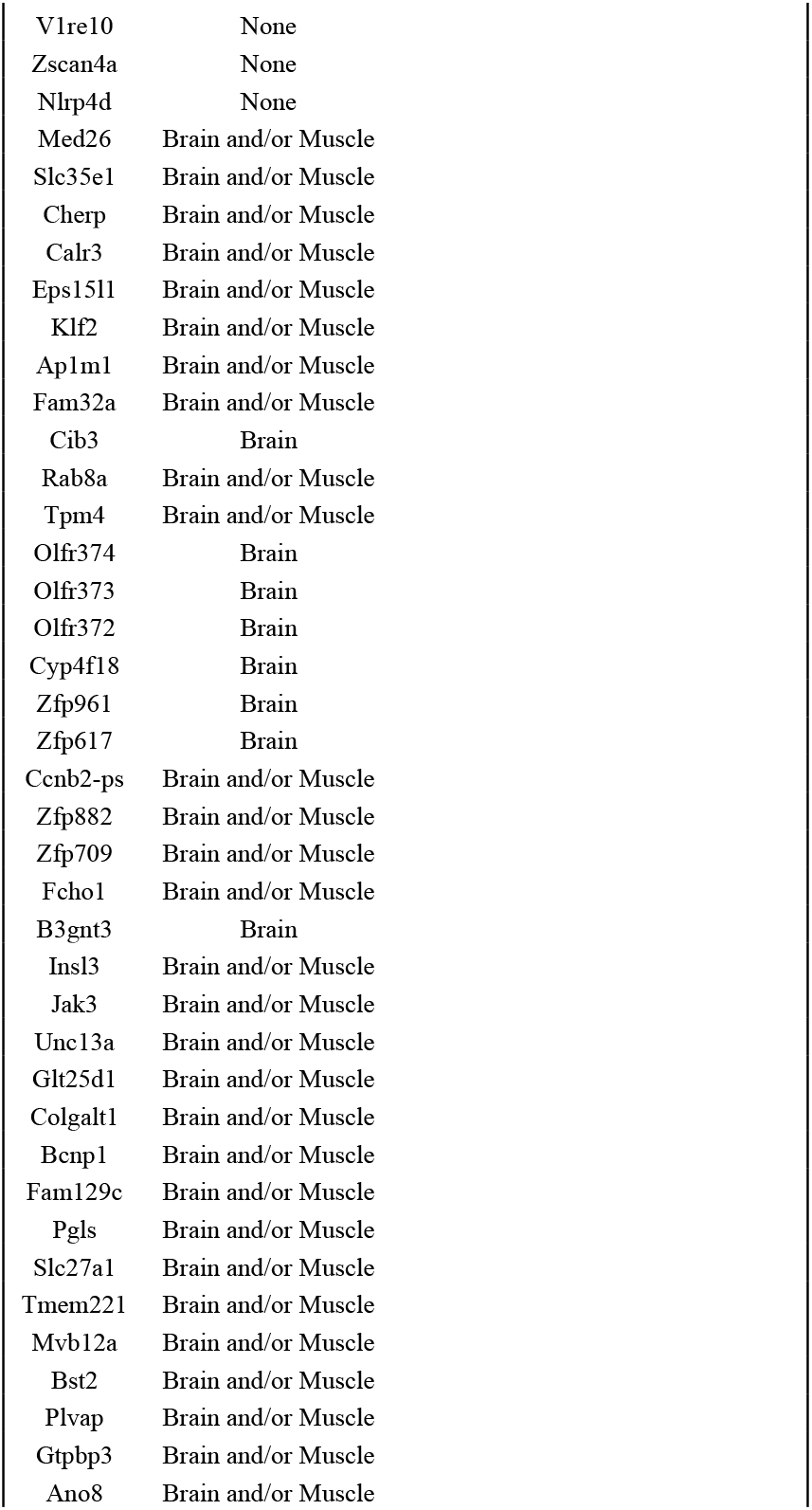
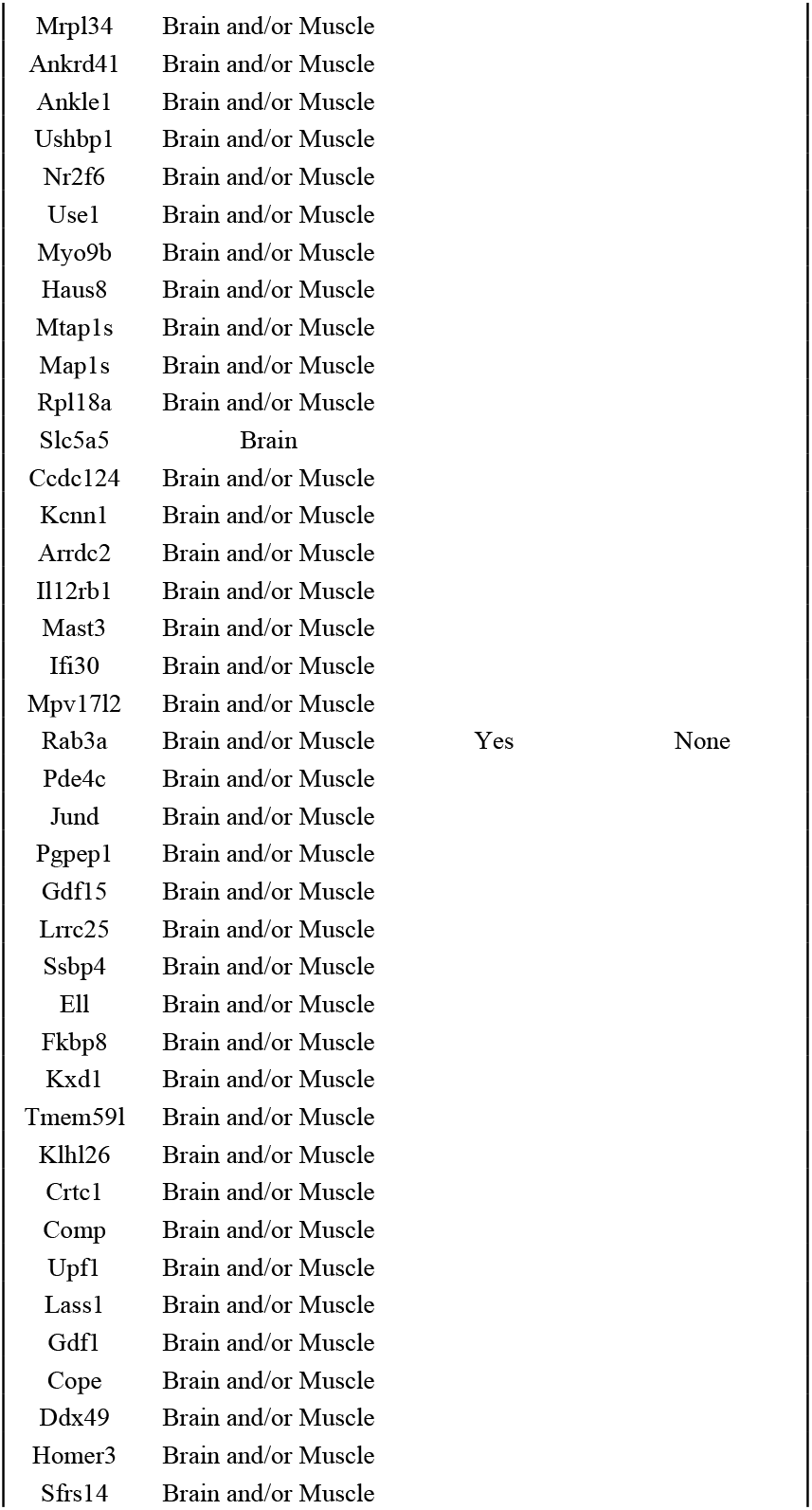
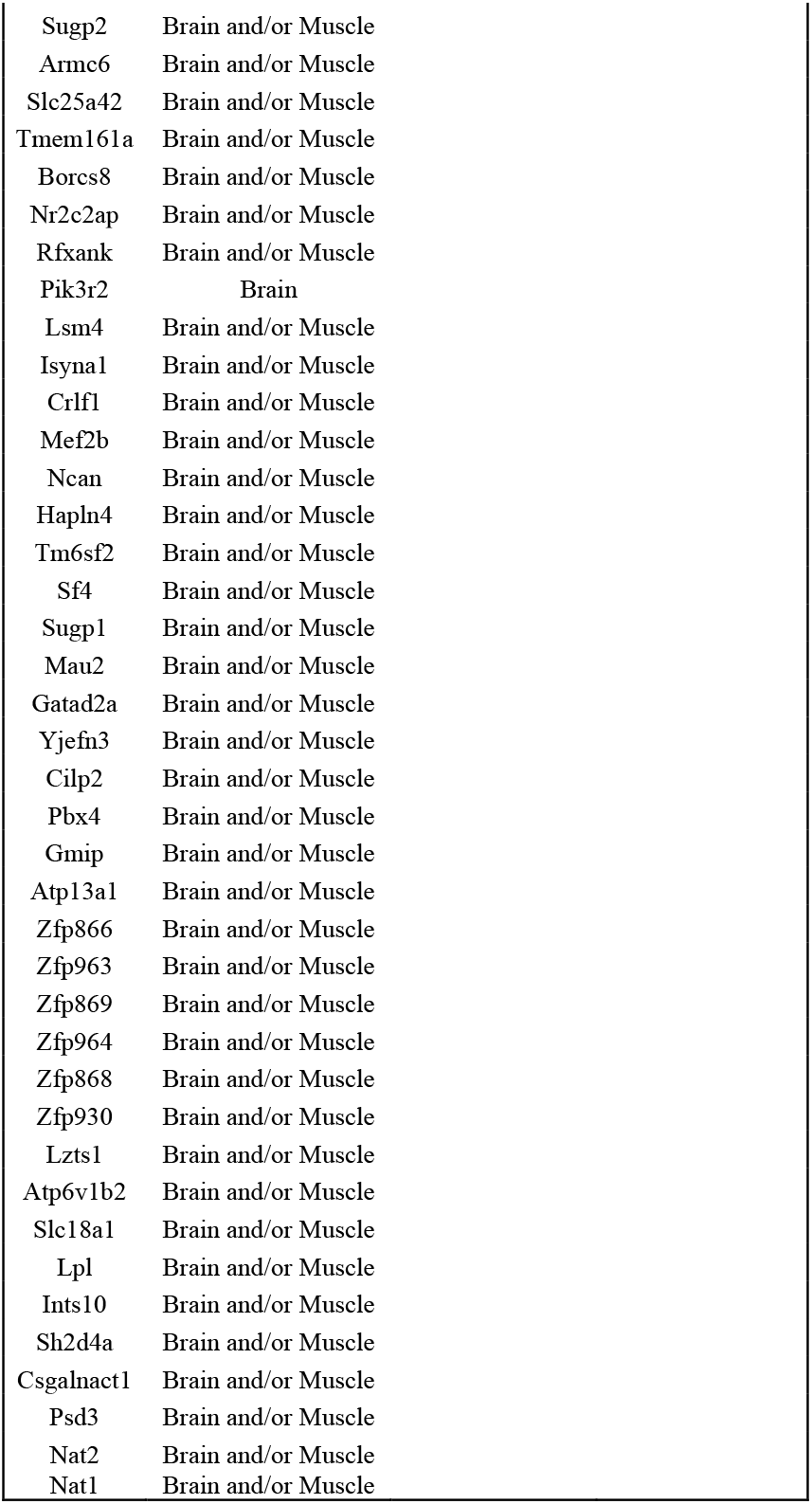
Identification of priority genes using three criteria. Empty areas show genes that did not meet the criterion system.

